# Cadherin 4 assembles a family of color-selective retinal circuits that respond to light offset

**DOI:** 10.1101/2024.05.08.593219

**Authors:** Aline Giselle Rangel Olguin, Pierre-Luc Rochon, Catherine Theriault, Thomas Brown, Michel Cayouette, Erik P. Cook, Arjun Krishnaswamy

**Affiliations:** Department of Physiology, McGill University, Montreal, Canada; Institut de Recherches Cliniques de Montréal, Montreal, Canada; Department of Anatomy and Cell Biology, McGill University, Montreal, Canada; Department of Medicine, Université de Montréal, Montreal, Canada

## Abstract

Retinal interneurons and projection neurons (retinal ganglion cells, RGCs) connect in specific combinations in a specialized neuropil called the inner plexiform layer (IPL). The IPL is divided into multiple sublaminae, with neurites of each neuronal type confined to one or a few layers. This laminar specificity is a major determinant of circuit specificity and circuit function. Using a combination of approaches we show that RGCs targeting IPL sublamina 1 and 3a express the adhesion molecule cadherin 4 (Cdh4). Using calcium imaging and iterative immunostaining, we classified Cdh4-RGCs into 9 types that each encode unique aspects of dark visual stimuli. Cdh4 loss selectively disrupted the layer-targeting of these RGCs, reduced their synaptic inputs from interneurons, and severely altered their visual responses. Overexpression of Cdh4 in other retinal neurons directed their neurites to s1-3a through homophilic interactions. Taken together, these results demonstrate that Cdh4 is a novel layer targeting system for nearly a third of all RGC.

## Introduction

The mouse retina is composed of ∼130 neural cell types that connect in specific and stereotyped ways to create circuits that encode unique aspects of the visual scene^1–4^. During development, amacrine (AC), bipolar (BC) and retinal ganglion cell (RGC) types restrict their axons or dendrites to one or two of the 5 sublaminae in the retina’s primary neuropil, the inner plexiform layer (IPL). This laminar targeting promotes synapses among co-laminar AC, BC, and RGC types and endows the latter with selective responsiveness to specific visual features. Yet, few of the molecules that direct axons and dendrites to each IPL sublamina are known. We previously showed an important role for some Cadherin isoforms (Cdhs) in the layer targeting RGCs and BCs involved in direction selective circuits^5,6^. This raised the possibility that Cdhs play a similar role in the assembly of other retinal circuits.

The expression of Cdhs in the retina is well documented in publicly available sequencing atlases^7–10^, yet few Cdhs have been studied for their roles in the laminar targeting of developing retinal neurons, particularly in RGCs. Our bioinformatic analysis identified several Cdhs that exhibited RGC subtype-specific expression patterns (**Fig.1A**). One of these, Cdh4, was discovered and originally named Retina- or R-cad by Takeichi and colleagues over 30 years ago^11^. Prior work on Cdh4 showed that it binds homophilically in vitro and is expressed by subsets of retinal interneurons and neurons in the ganglion cell layer^12–15^. These results motivated us to ask if Cdh4 acts as a laminar targeting molecule.

**Figure 1.**
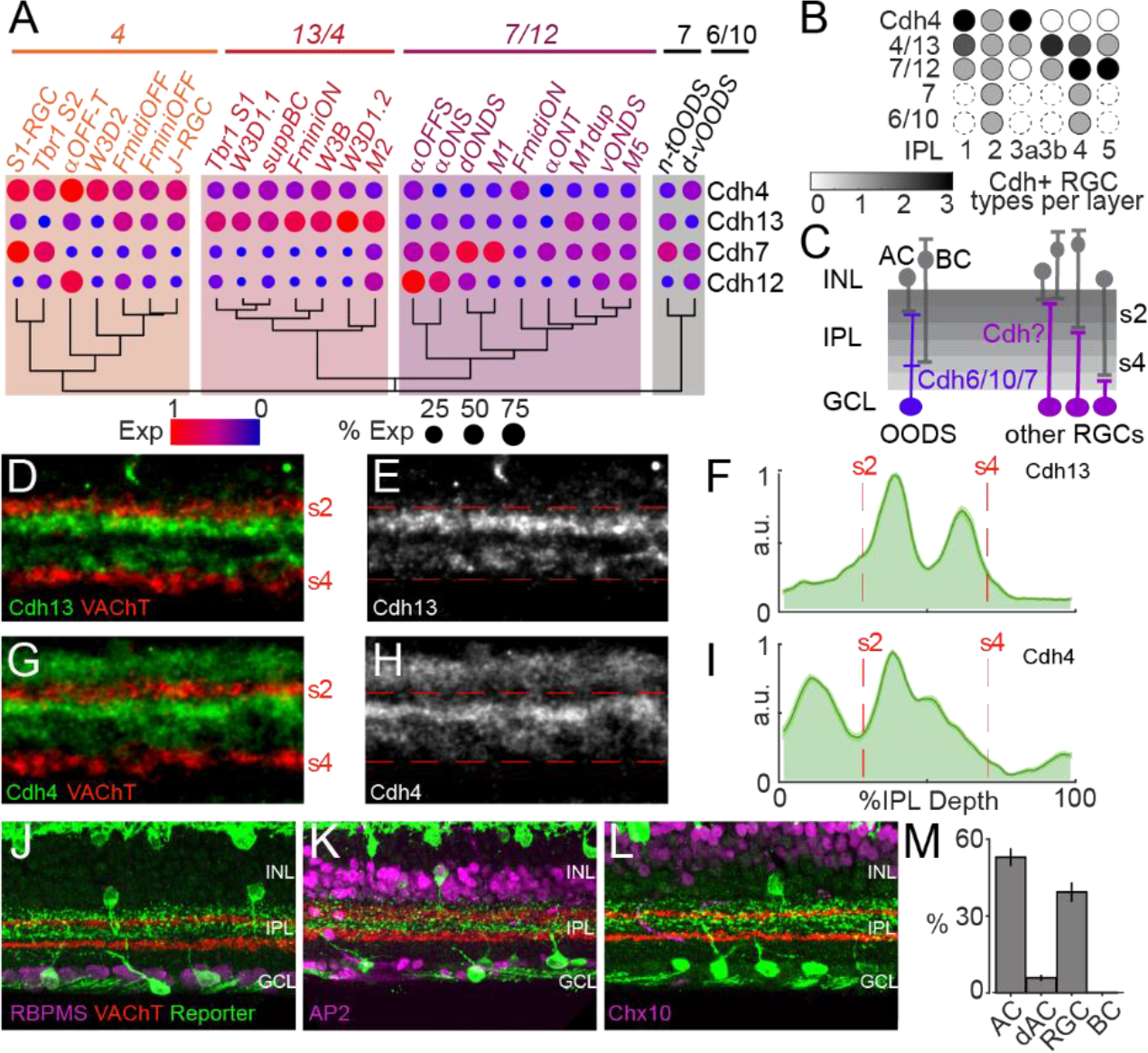
RGCs projecting to the same IPL sublamina express a common Cdh. A. Dendrogram of known RGCs based on their differentially expressed Cdhs. Hierarchical clustering organizes known monostratified RGC types by expression of a unique combination of Cdhs 4, 7, 12, and 13. Expression of Cdhs6&10 in v- and t-ooDSGCs from refs 5-6 also shown. **B.** Approximate dendritic stratification of RGCs in A. Cdh4 labels RGCs that project to s1 and s3a, Cdh4/13 RGCs project to s3a&b, and Cdh7/12 RGCs project to s4-5. **C.** Cartoon of IPL layers with known Cdh-RGC- sublayer associations indicated. Other RGCs may express different Cdhs to find appropriate sublayers. **D-I**. Retinal cross-sections stained with antibodies against vesicular acetylcholine transporter (VAChT), Cdh13 (D-E), Cdh4 (G-H), and average line scans (±SEM) taken through all such experiments (F-I, n=4/condition). **J-L**. Retinal cross-sections from a Cdh4-Het mouse crossed to a Cre-dependent reporter stained with markers for RGCs (RBPMS, J), ACs (Ap2α, K), and BCs (Chx10, L). **M.** Percent of amacrine (AC), displaced amacrine (dAC), retinal ganglion (RGC), and bipolar (BC) cells expressing Cdh4, quantified from experiments like J-L (n = 4 mice).

We found Cdh4 expression within a subset of RGCs and ACs targeting IPL sublamina 1 and 3a. Using genetic tools and an iterative immunostaining method, we dissected Cdh4-RGCs into 7 known and 2 predicted novel RGC types. Two-photon calcium imaging showed that Cdh4-RGCs detected various features of dark visual stimuli which included preferences for color. Loss of Cdh4 led the dendrites of most of these RGCs to grow aberrantly into inappropriate lamina, disrupting their synaptic inputs and impairing their visual responses. Moreover, when Cdh4 was overexpressed in ACs and BCs, these neurons targeted their neurites into sublamina 1 and 3a by a homophillic mechanism. Taken together, our results show that Cdh4 is a novel layer-targeting system for nearly a third of all RGCs . More generally, these and our previous results^5,6^ suggest that a cadherin code plays a major role in determining laminar specificity throughout the IPL and raise the possibility that similar codes could be involved in laminar specificity elsewhere in the central nervous system.

## Results

### Retinal neurons projecting to a unique sublamina express unique Cdhs

To investigate whether Cdhs guide axons and dendrites of retinal neurons to specific IPL sublaminae, we examined patterns of Cdh expression within publicly available RGC sequencing atlases^7–10^. By comparing Cdh expression across all RGC types we identified the most differentially expressed Cdh members and related their expression pattern to RGC types growing their dendrites within a single IPL layer. This analysis revealed a set of 4 Cdhs expressed by RGCs with unique IPL layering patterns (**Fig. 1A**).

Some of the RGC types expressing high levels of Cdh4 have been reported to have dendrites in sublamina 1&3a (s1&3a)^9,16–21^, whereas those that express the other 3 Cdhs have been reported to target different layers. RGC types with high levels of Cdh13 are reported to target s3b^16,17,19^; and those with high Cdh7&12 are reported to target s4&5 (**Fig. 1B**)^16,17^. These results raise the possibility that each RGC group uses its respective Cdh to find these layers (**Fig. 1C**).

To investigate this idea further, we stained retinal cross-sections with antibodies against either Cdh4 or Cdh13 and found that Cdh4 and Cdh13 labelled unique IPL sublaminae (**Fig. 1D-I**). Cdh13 stained a pair of bands in s3a and s3b (**Fig. 1D-F**), whereas Cdh4 stained a pair of bands in s1 and s3a (**Fig. 1G-I**). Antibodies to vesicular acetylcholine transporter (VAChT) marked s2 and s4^5,6^ ^20,21^. (We were unable to find specific antibodies for Cdhs7&12.) These results suggest that Cdhs4&13 could target RGC dendrites to specific IPL sublaminae.

### Cdh4 labels ACs and RGCs projecting to IPL sublaminae 1 and 3a

We focused on Cdh4 for two reasons. First, it was the only Cdh in our analysis that exclusively marked a set of RGC types; all other Cdhs appeared in combination. Second, the RGC types predicted to exclusively express Cdh4 are among the most numerous^9^ and Cdh4 antibody labelling showed a pair of thin strata (**Fig. 1G-I**), facilitating analysis of potential loss-of-function phenotypes.

To investigate the role of Cdh4 in laminar targeting, we used mice in which the Cdh4 gene is interrupted by a construct encoding a Cre-recombinase-human estrogen receptor fusion (CreER)^16^. As heterozygotes (Cdh4-Het), Cdh4-CreER lets one mark and manipulate Cdh4-exressing cells. As homozygotes this line is also a Cdh4 null (Cdh4-KO). Crossing Cdh4-Hets to Cre-dependent reporter lines and treating the progeny with tamoxifen revealed many labelled neurons in the Ganglion Cell Layer (GCL) that co-expressed the RGC marker RBPMS (**Fig. 1J&M**). We also found a large population of neurons in the inner nuclear layer that co-expressed the AC marker AP2α (**Fig. 1K&M**). While we saw neurons whose position and anatomy matched those of horizontal cells, we never saw reporter-labelled neurons that co-expressed the BC marker Chx10 (**Fig. 1L-M**). We were, however, able to sparsely label some Cdh4+ve ACs in wholemounts and quantify their lamination in rotated z-stacks (**Supplemental Fig. 1A-C**). Most of these Cdh4-ACs (**Supplemental Fig. 1D-E**) innervated s1-s3a, as expected from the bulk labeling pattern in **Fig. 1K**. These results indicate that a subset of RGCs and ACs express Cdh4 on their dendrites which together concentrate in IPL s1 and s3a.

### Cdh4 defines a family of 9 RGC types

Our bioinformatic analysis predicted that at least 9 transcriptomically defined RGC types express high levels of Cdh4 (**Fig. 1A**). Atlas annotation suggested that seven of these clusters were (i) alpha RGCs with transient responses to light offset (αOFF-T)^17^, (ii) color-opponent RGCs that show orientation- and motion preferences (J-RGCs)^20–22^, (iii) S1-RGCs named for their expression of Sdk1^23^, (iv) an RGC that expresses Tbr1 (Tbr1_S2)^18^, (v-vi) two RGC types expressing Foxp2 (Fmini-OFF, Fmidi-OFF)^16^, (vii) and a poorly characterized narrow-field RGC (W3D2)^19,24,25^. Two other clusters were predicted to be novel members of the Tbr1+ and Foxp2+ RGC family (T-Novel and F-Novel)^9^.

We defined a set of marker genes that could combinatorially label each Cdh4-RGC type (**Fig. 2A**). The integral membrane protein Tusc5/Trarg1 was an important part of our marker definitions, but we could not find specific antibodies against this protein. To circumvent this, we took advantage of the observation that all Tusc5+ve RGCs are labelled with YFP in the TYW3 transgenic line^9^. Briefly, TYW3 exhibits insertion-site-dependent expression of YFP in several RGC types that grow dendrites into unique regions of s1-3b. These include the W3D2 (s3), FminiOFF (s1), and Tbr1_S2 RGCs (lower s1, upper s2)^9^. Thus, we reasoned that a mouse carrying one copy of Cdh4-CreER (Cdh4-Hets), Cre-dependent TdTomato (Ai27D), and the TYW3 transgene could be used to distinguish the Cdh4-RGC types. Treating such triple- genotype Cdh4-Hets with tamoxifen labelled all Cdh4-RGCs with Tdtomato, but a subset of these cells also constitutively expressed YFP (TYW3). To distinguish each Cdh4-RGC type, we stained retinal cross-sections from these mice with antibodies against the transcription factors Tbr1, Brn3c, and Foxp2, the cytokine Osteopontin (Ost), YFP, and Tdtomato (**Fig. 2A**).

**Figure 2.**
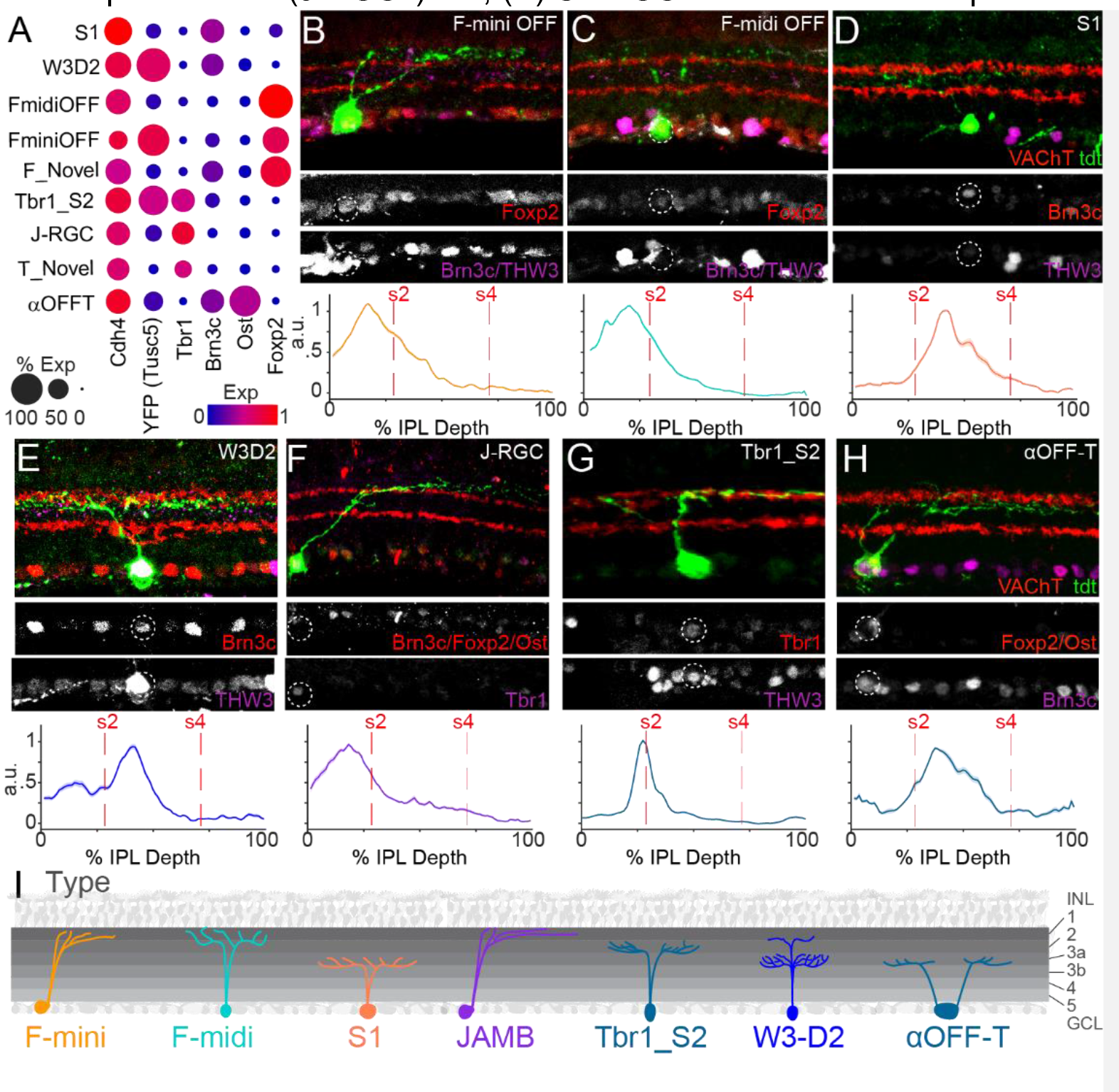
A family of Cdh4 RGCs. A. Dot plot showing the expression of Cdh4, Tusc5 (TYW3), Tbr1, Brn3c, Osteopontin (Ost), and Foxp2 across Cdh4-RGC types. **B-H. Top**, retinal cross- sections taken from a Cdh4-Het/THWGA3 that also expresses Cre- dependent tdtomato (green). Sections were counterstained with Foxp2 / Brn3c / YFP to identify mini- and midiOFF RGCs (**B,C**); Brn3c / YFP to mark S1- and W3D2 RGCs (**D,E**); Brn3c / Foxp2 / Ost / YFP / Tbr1 to mark JAM-B and Tbr1-S2 RGCs (**F,G**) and Foxp2 / Ost / Brn3c to mark αOFF- T RGCs (H)**. Bottom,** Line scans (±SEM) taken through all experiments like those shown at top. N ≥ 6/RGC type from 18 mice. **I.** Cartoon of the 7 Cdh4-RGCs. Novel types not shown.

Fmini- and Fmidi-OFF RGCs co-expressed Foxp2 but were distinguished from each other because only FminiOFFs were YFP+ve (**Fig. 2B-C**). Both F-RGCs grew dendrites into s1 of the IPL. S1- and W3D2 RGCs expressed Brn3c but only W3D2 expressed YFP (**Fig. 2D-E**). S1-RGCs targeted the interface between s3a and s3b, whereas W3D2 grew dendrites in s1&s3a. J- and Tbr1_S2 RGCs co-expressed Tbr1+ but only Tbr1_S2s were YFP+ (**Fig. 2F-G**). Both J- and Tbr1_S2s grew thin dendritic arbors that targeted the lowest part of s1 but the latter’s dendrites grew near the s1-s2 interface. Finally, αOFF-T expressed Ost+/Brn3c+ and grew dendrites in s3a (**Fig. 2H**). Some Foxp2+/YFP-ve and Tbr1+/YFP-ve RGCs did not match the anatomy FmidiOFF and J-RGCs (**Supp. Fig. 2A-B**), suggesting that these RGCs are likely to be the ‘novel’ F- and T-RGC types that express Cdh4 (**Fig. 2A**).

Simultaneous labeling of retina with many of our Cdh4-RGC markers (**Fig.2A**) was not feasible because many of these antibodies were raised in the same species. Therefore, to assess the completeness of our Cdh4-RGC molecular taxonomy, we adapted iterative indirect immunofluorescence imaging methods (4i) ^26,27^; this allowed us to serial stain retinae with multiple antibodies (**Supp. Fig. 3D**). Using 4i, we re-stained wholemount retinae from Cdh4-Hets bearing THWGA, and Cre-dependent Tdtomato transgenes with all our Cdh4-RGC markers (**Fig.2A**) and quantified the proportion of reporter+ve/marker-ve cells (**Supp. Fig. 2C**). We saw a negligible number of marker-ve neurons, indicating that our molecular taxonomy of Cdh4-RGC types (**Fig.2A**) is likely complete. This analysis also demonstrates that the Cdh13-RGCs from our bioinformatic survey are Cdh4 negative. Thus, a family of 7 known and 2 novel RGC types expresses Cdh4 but not Cdh13 and targets s1 and/or s3a (**Fig. 2I**).

### Cdh4-RGCs are strongly driven by light offset and show preferences for color

Laminar organization in the IPL is a major determinant of visual feature selectivity^2,3^. For example, BCs that sense light offset (OFF) innervate s1-s3a whereas those sensing light onset (ON) innervate s3b- 5^28,29^ respectively. RGCs with dendrites in one of these layers inherit the ON or OFF sensitivity of their BC inputs which divides RGCs into three families: OFF (s1-s3a), ON (s3b-s5), or ON-OFF types^3^. Given that Cdh4 labels RGCs that stratify in s1-s3a (**Figs1-2**), we predicted that Cdh4-RGCs would exhibit strong OFF-responses.

To characterize the visual responses of Cdh4-RGCs, we crossed Cdh4-Hets to Cre-dependent GCaMP6f lines and used two-photon microscopy to record RGC responses in retinal explants to a set of stimuli that measure feature selectivity^23,30^ (**Fig. 3A-C**). Cdh4-RGCs exhibited diverse responses to these stimuli (**Fig. 3D**) which included (i) full-field flashes that varied with temporal frequency (t-chirp), (ii) contrast (c- chirp), (iii) or color (green/blue).

**Figure 3.**
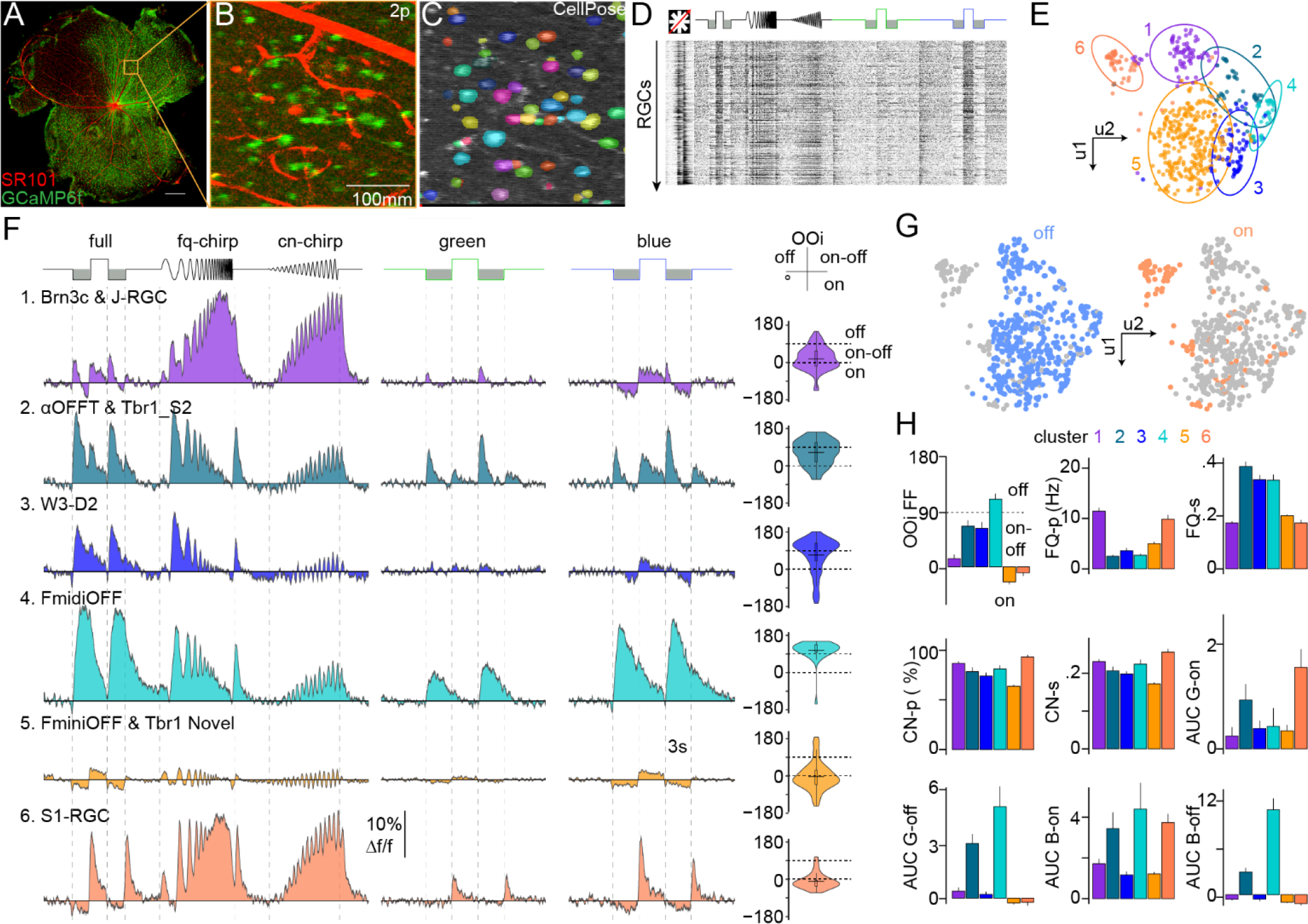
Cdh4 RGCs react strongly to light offset and exhibit preference for stimulus color. A-C. Low power image of a Cdh4-Het x Ai32 retina showing GCaMP6f+ve RGCs (green) and blood vessel staining using sulphorhodamine101 (SR101, red) (A). Inset magnifies a sample field (B) and shows ROIs detected by CellPose (C). **D.** Response matrix for 788 RGCs taken from imaging several fields like that shown in A-C. **E.** UMAP represents the responses of individual Cdh4-RGCs. Colors and ellipses obtained from GMM-based clustering in all 14 principal components. **F.** Mean response of each cluster shown in E to a full-field flash (full), a full-field flash that becomes faster every cycle (fq-chirp), a flash that grows in contrast every cycle (cn-chirp), a flash with only the green LED, and a flash with only the blue LED. Violin plots at right show the on-off index computed for cells belonging to each cluster with ranges for on, off, and on-off RGCs indicated. **G.** Umap plot colored according to whether an RGC has an On-Off index higher than 0 (Off) or lower than 0 (On). **H.** Bar graphs quantifying average on-off index from the full-field response (OOi), frequency preference and selectivity from fqchirp (FQ-p, FQ-s), contrast preference and selectivity from cn-chirp (CN-p, CN-s), and the integrated calcium response (area under the curve, AUC) to green (G) or blue (B) full-field flashes for the indicated transitions (Gon, G-off, B-on, B-off)

To analyze these responses, we reduced our dataset to 14 principal components that accounted for the most response variability (**Supp Fig. 3E-G**); we clustered responses using gaussian mixture models, and computed the cosine similarity of clustered traces to all cluster means to assess cluster purity (**Supp. Fig. 3H-I, Supp. Fig. 4A**). This analysis grouped Cdh4-RGCs into 6 functional clusters that we visualized with uniform manifold approximation (UMAP) (**Fig. 3E**). Each cluster showed a unique average response to our stimuli (**Fig. 3F**). Most clusters were strongly driven by light offset (**Fig. 3F-H**, *full*), indicating that they are OFF RGCs. One cluster showed ON response full-field responses which matches the S1-RGC^23^ (see below). All clusters preferred high contrast but showed unique temporal frequency preference (**Fig. 3F**, *fq-chirp*, *cn-chirp*).

Most clusters showed various forms of color sensitivity. For example, clusters 2,4, and 6 showed weaker responses to green stimuli than to blue (near UV) stimuli (**Fig. 3F**, *green, blue*). On the other hand, clusters 1-3 showed OFF responses to green flashes but ON and OFF responses to blue flashes (**Fig. 3F***, green, blue*), suggesting that they contained color-opponent RGCs. Comparing responses across clusters provided a measure of the RGCs’ relative preferences for the other stimuli (**Fig. 3H**).

These results indicate that Cdh4-RGCs encode light offset, or dark visual stimuli, and show preferences for stimulus color. Since YFP from the THWGA3 transgene is required to distinguish J-RGCs from Tbr_S2s, FminiOFFs from FmidiOFFs, and W3D2s from S1-RGCs, we could only perform a partial match of our molecular (**Fig.2A**) and functional Cdh4-RGC definitions (**Fig.3E**). Briefly, we 4i stained retinae following calcium imaging, related Cdh4-RGC marker expression to functional response, and then used prior descriptions of the Cdh4-RGC types to assign them to clusters.

Brn3c, Tbr1, and Foxp2 labelled pairs of clusters (**Supp. Fig 4B**), consistent with the expression of these markers across Cdh4- RGCs: Brn3c marks W3D2 & S1-RGC (*Clusters3&6*); Tbr1 marks J-RGC and Tbr1_S2 (*Clusters1&2*); and Foxp2 marks Fmini and FmidiOFFs (*Clusters4&5*) (**Fig.2A, Supp. Fig 4B**).

Cluster 2 had high levels of Ost, matching descriptions of αOFF-Ts^17^. Cluster 1 and cluster 2 also had high levels of Tbr1, suggesting the presence of J-RGCs and Tbr1_S2s^18^ (**Supp. Fig 4B**). However, the high temporal frequency preferences of cluster 1 matched that of J-RGCs^30^ (**Fig.3F)**. Given this, we placed J-RGCs in cluster 1 and Tbr1_S2 in cluster 2.

Clusters 3&6 had high Brn3c (**Supp. Fig 4B**), but cluster 6 showed the strong full-field On-responses that characterize the S1- RGC^23^. We thus inferred that S1-RGCs were in cluster 6 and W3D2s were in cluster 3. Finally, clusters 4&5 showed high Foxp2, but cluster 5 contained the most RGCs in our dataset which is consistent with the FminiOFF density that is the highest of all Cdh4-RGCs^9,16^. For this reason, we inferred that FminiOFFs were in cluster 5 and FmidiOFFs were in cluster 4. Taken together, our data show new molecular and functional definitions for six OFF- and color-sensitive, Cdh4-RGC types.

### RGC dendrites require Cdh4 to target appropriate IPL lamina

Sensitivity to light offset^28^ and green stimuli^31^ are features that have been linked to BCs and ACs whose processes innervate the upper half of the IPL. The correspondence of these properties with those of Cdh4- RGCs led us to test whether Cdh4 was required for their dendritic lamination. To this end, we compared overall lamination in reporter-labelled cross-sections from Cdh4-Hets to those of Cdh4-KOs which bear two copies of Cre-ER that interrupt Cdh4 gene expression^16^.

Loss of Cdh4 produced no noticeable change to the gross organization of the cellular and neuropil layers in the retina (**Supp. Fig. 5A-F**). In contrast, we saw striking mistargeting phenotypes in reporter- positive dendrites in Cdh4-KO retinae that included ectopic projections in sublaminae 4 and 5 (**Fig. 4A**). These data show that a subset of retinal neurons (ACs and RGCs) require Cdh4 to enter s1 and s3a.

**Figure 4.**
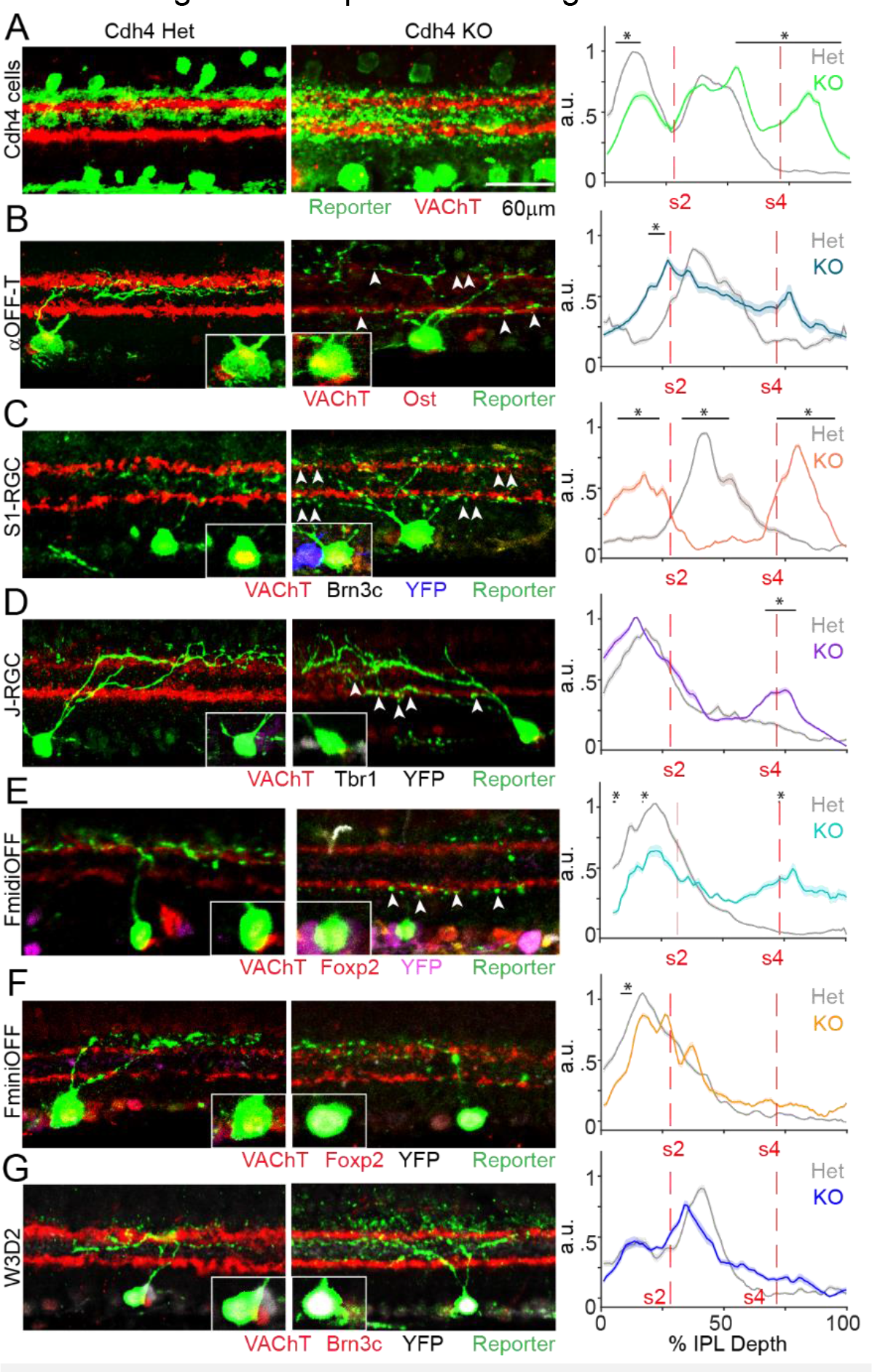
Loss of laminar targeting in the absence of Cdh4. A. Cdh4-Het control (left) and Cdh4-KO (middle) retinal crossections stained for antibodies against reporter and VAChT. Right: Linescans taken through experiments like those shown on the left and middle. **B-G.** Cross sections showing αOFF-T (B), S1 (C), J (D), FmidiOFF (E), FminiOFF (F) and W3D2 (G) RGCs labelled with Tdtomato in retinae from Cdh4-Het (left) and Cdh4-KO (middle) mice crossed to TYW3. Triangles indicate ectopic branches in Cdh4-KOs. Each section was stained with a panel of type-specific antibodies. Right, mean intensity linescans show mean ± SEM fluorescence across the IPL divided into 88 bins. N ≥ 6 for each RGC from more than 12 mice/condition. * = P < 0.05 t-test, for the indicated positions.

The laminar deficits seen in our bulk labelling (**Fig. 4A**) might reflect deficits on all Cdh4-RGCs and ACs or just a subset. To better resolve the RGCs, we delivered retrogradely infecting AAVs to the visual thalamus (LGN) of Cdh4-Hets and Cdh4-KOs who also bore the THWGA3 transgene. This procedure sparsely labelled the αOFF-T, S1-RGC, J-RGC, FminiOFF, FmidiOFF, and W3D2; Tbr1_S2 were rarely labelled, suggesting that they do not innervate the LGN. The loss of Cdh4 severely impacted αOFFT- and S1-RGCs who showed dendritic strata within the ON-laminae (**Fig. 4B-C**). J-RGCs and FmidiOFFs in Cdh4-KOs often exhibited ectopic branches that grew from their primary dendrite into sublamina 4-5 (**Fig. 4D-E)**. Dendritic deficits on FminiOFF and W3D2 were modest (**Fig. 4F-G**).

As a control against non-specific targeting deficits in the absence of Cdh4, we bred Cdh4-KOs to a transgenic line (HB9-GFP) that fluorescently labels DS-RGCs expressing Cdh6/10^5,6,32^ and examined the morphology of this cell in retinal sections. This analysis showed that the dendritic lamination of DS-RGCs in Cdh4 nulls were like those in controls (**Supp. Fig. 4C-D**). Taken together, these results show that Cdh4 is required for the dendritic targeting of several OFF-RGC types.

### Loss of Cdh4 diminishes OFF responses and color selectivity

We next asked whether Cdh4-RGC exhibit functional deficits in the absence of Cdh4. To do this, we imaged the responses of these RGCs in Cdh4-KOs to the same stimulus set as we used in controls and projected each RGC response vector into the 14 components obtained from our control dataset (*see methods*). We then computed the Euclidean distance from each Cdh4-KO RGC to the center of every control cluster and assigned such RGCs to the nearest control cluster (**Fig. 5A-B**, **Supp. Fig. 3F-J**, *see methods*). In general, the proportion of RGCs in control and Cdh4-KO clusters were similar (**Fig. 5C**). However, the average visual responses were significantly diminished by Cdh4 loss (**Fig. 5D**).

**Figure 5.**
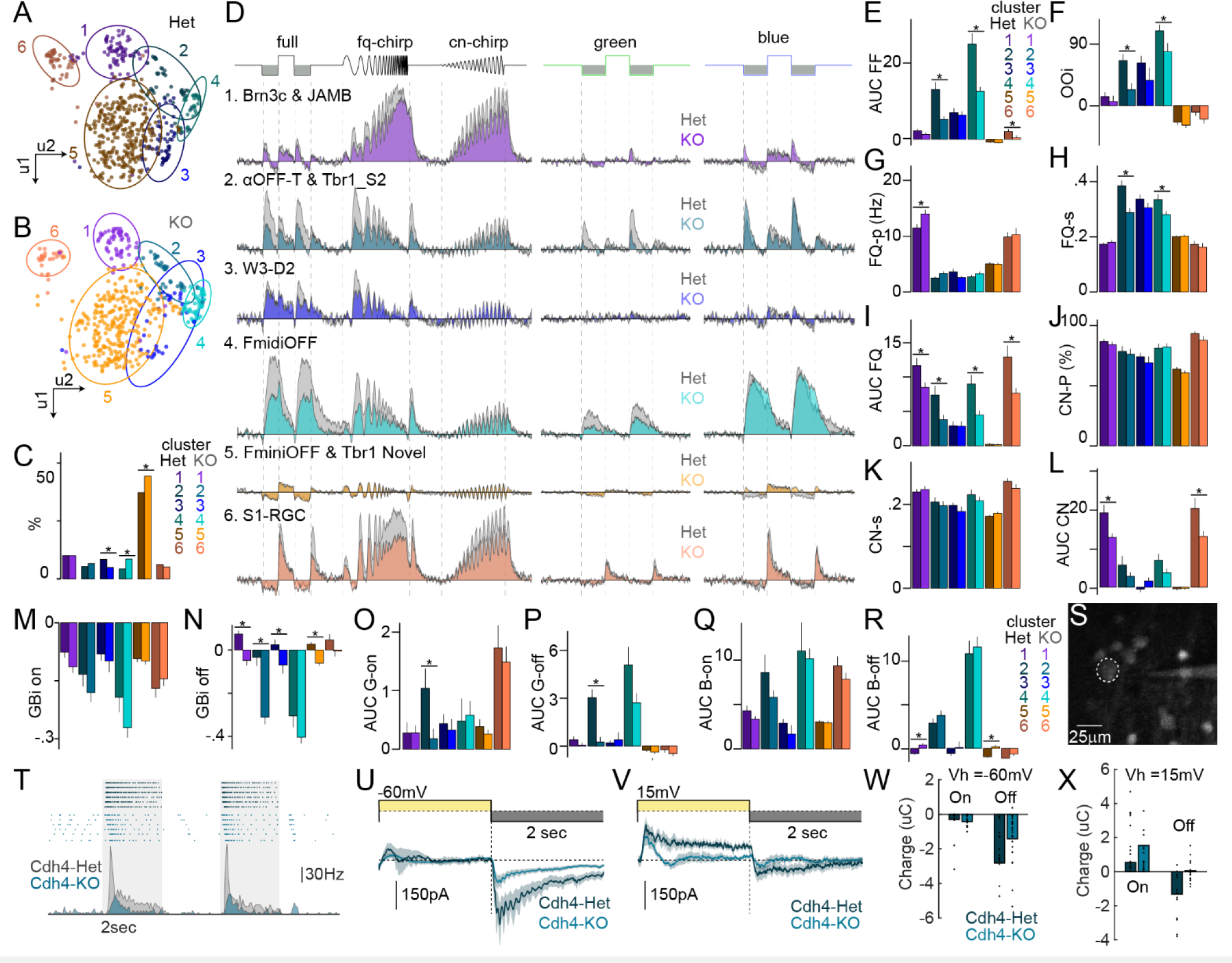
Cdh4 loss impairs responses to light offset and to color. A-B. UMAPs for Cdh4-RGCs with clusters indicated for recordings in Cdh4-Het controls (A) and Cdh4-KO retinae(B). **C.** Bar graph showing the percent of RGCs in control (n = 788 cells in 7 mice) and Cdh4 null (n = 611 cells in 6 mice) clusters as shown in A-B. **D.** Mean response of each cluster shown in A-B to a full-field flash (full), a full-field flash that becomes faster every cycle (fq-chirp), a flash grows in contrast every cycle (cn-chirp), and a flash with only the green or blue LED. Average control responses shown in gray and Cdh4-KO in color. **E.** Integrated calcium response (area under the curve, AUC) of the calcium response to a full field flash for control and Cdh4-KO clusters. **F.** Average On-Off index value for control and Cdh4-KO clusters. **G-I.** Preferred stimulus frequency (G), frequency selectivity (H), and AUC to fqchirp stimulus (I) for control and Cdh4-KO clusters. **J-L.** Preferred stimulus contrast (J), contrast selectivity (K), and AUC for cn-chirp stimulus (L) for control and Cdh4-KO clusters. **M-R.** Bar graphs comparing the average Green-Blue index value (GBi on, M; GBi off, N) and AUC for green (G) or blue (B) full-field ON or OFF stimuli (Gon, O; G-off, P; B-on, Q; B-off, R) for control and Cdh4-KO clusters. **S.** Example two-photon targeted recording from a control αOFFT-RGC (S) **T,**Top: raster of responses to a full-field flash in a control and a Cdh4-KOs αOFFTRGC. Bottom: Average spike rate of responses like those showed on top. **U.** Average ON and OFF evoked currents recorded at Vh = -60 mV in αOFFT-RGCs. **V.** Average ON and OFF evoked currents recorded at Vh = +15 mV in αOFFT-RGCs. **W-X**. Bar graphs show average peak response for currents shown in U-V. * = P < 0.05 reported from a t-test between the conditions.

OFF responses to the full field flash were strongly attenuated in clusters 2&4 with smaller reductions in their ON-responses (**Fig. 5D-E**, *full*). This led to a significantly lowered selectivity for light offset (**Fig. 5F**). In addition, most clusters had reduced responses to frequency and contrast chirp stimuli (**Fig. 5D, I, L**), but their preference and selectivity for these stimuli were unchanged (**Fig. 5G-H, J-K**). Selectivity for green/blue OFF-stimuli was significantly reduced with little change in green/blue ON-stimuli (**Fig. 5M-N**), consistent with the overall preference for light offset across Cdh4-RGCs. Responses to green flashes were more impaired than responses to blue ones (**Fig. 5O-R**), suggesting that mistargeted Cdh4-KO RGCs cannot connect with ACs and/or BCs carrying signals originating in green cones or possibly rods^20^. Finally, we observed that responses from clusters 3&5 were more resistant to Cdh4-loss (**Fig. 5D-R**), consistent with our anatomical studies showing that FminiOFFs & W3D2s made fewer dendritic targeting errors. To control against a general effect of Cdh4 loss on RGC light responses, we also examined the calcium responses of all DS-RGCs in Cdh4-KO retinae (*see methods*) and found direction selective responses that were similar in amplitude and selectivity to those found in controls (**Supp. Fig.4E**). Taken together, these studies indicate that Cdh4 loss impairs the light responses of most Cdh4-RGCs.

We predicted that the functional deficits in Cdh4-KOs RGCs arose because their mistargeted dendrites could not synapse with appropriate inputs. To test this idea, we patched reporter+ve αOFF-Ts (identified by their large somata) (**Fig. 5S**) to measure excitatory (BC) and inhibitory (AC) synaptic currents on these neurons. Loose-patch recordings from control αOFF-Ts showed the OFF-evoked transient spiking characteristic of αOFF-Ts (**Fig. 5T**). Recordings from Cdh4-KO αOFF-Ts showed significantly weaker spike responses. Whole-cell recordings in control αOFF-Ts held at negative holding potentials (-60mV) showed small, outward currents to light onset and large inward currents to light offset (**Fig. 5U**). Recordings at positive holding potentials (+15mV) showed large outward currents to light onset and weak inward currents to light offset (**Fig. 5V**). The same recordings from αOFF-Ts in Cdh4-KOs showed significant loss of OFF-inward responses at -60mV and ON-outward responses at +15mV (**Fig. 5W-X**). These results indicate that Cdh4 loss reduces functional synapses between αOFF-Ts and both BCs and ACs. Taken together, we conclude that Cdh4 loss selectively impairs Cdh4-RGCs visual responses, likely from a loss of synaptic input.

### Cdh4 misexpression directs neurites to s1-3a

We next asked whether Cdh4 acts homophilically to target neuronal processes to s1 and 3a. To test this idea, we misexpressed Cdh4 in developing retinal neurons with electroporation methods and asked if this directed neurites into sublamina 1 and 3a. Briefly, we obtained a cDNA construct encoding full-length Cdh4 with a C-terminal OFP fusion and validated its surface expression in heterologous cells using Cdh4 antibodies (**Supp. Fig. 5G**). Next, we electroporated Cdh4-OFP or a control reporter construct into the eyes of ∼P0 wild type mice and examined the morphology of labelled neurons in retinal cross-sections two weeks later.

We saw enrichment of neural processes in s1-s3a in retinae electroporated with Cdh4-OFP versus those electroporated with reporter (**Fig. 6A&C**). This result suggested that Cdh4- overexpressing neurites bind to endogenous Cdh4 present on the processes of Cdh4-RGCs and ACs s1-s3a. To test this idea, we electroporated Cdh4- OFP into the retinae of Cdh4- KOs and saw attenuated targeting (**Fig 6B&D**). These results indicate that processes ectopically expressing Cdh4 require an endogenous Cdh4 target to direct lamination. This result differed from our prior work with ectopically expressed Cdh8&9 which directed targeting without endogenous expression (see discussion).

**Figure 6.**
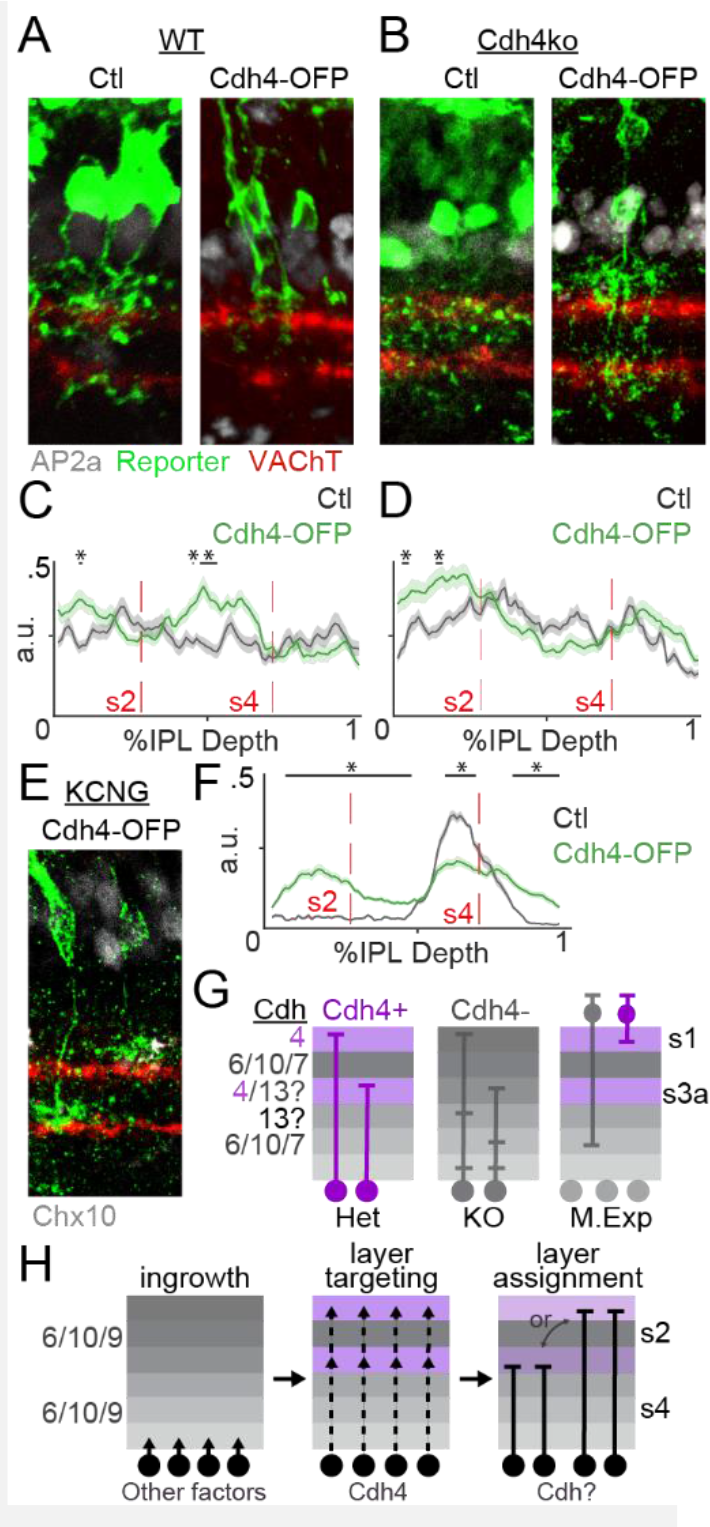
Cdh4 misexpression redirects retinal processes to s1-3a. A. Example retinal cross sections taken from wildtype retinae electroporated with reporter (Ctl) or Cdh4-OFP (A) constructs. **B.** Example retinal cross sections taken from Cdh4-KO retinae electroporated with reporter (Ctl) or Cdh4- OFP constructs. C, **D.** Line scans from all experiments like those as those shown in A,B. **E,** Retinal cross section from a KCNG-Cre mouse electroporated with constructs encoding Cre-dependent Cdh4- OFP. **F.** Linescans from all experiments like those shown in E. Gray linescan is a re-analysis of previously published control electroporations in KCNG retinae [**ref.5**]. **G.** Cartoon showing known Cdh to IPL layer associations. **H.** Speculative Cdh-based wiring model with three stages of retinal development. Following dendrite polarization and initial ingrowth, a combination of Cdhs would guide neurons to pairs of layers and assign them to specific layers. * = P < 0.05 t-test, for the indicated positions and N ≥ 15 sections and 4 mice for each condition. Linescans measure the IPL in 88 bins and are mean ± SEM fluorescence

To better resolve Cdh4’s effects on laminar growth, we repeated our electroporation with constructs containing Cre-dependent Cdh4-OFP or Cre-dependent reporters in neonatal KCNG4-Cre mice which grant genetic access to Type5 BCs. These BCs normally target sublamina 4^5^, but overexpression of Cdh4 in these neurons led them to form new branches in sublamina 1(**Fig. 6E-F**). Taken together, these results demonstrate that homophilic Cdh4-mediated interactions target neurites to s1-3a.

## Discussion

Here, we investigated whether the laminar choices of RGCs are directed by expression of Cdh4. Our study began with bioinformatic screens that revealed Cdh4 expression within a large group of RGCs and a subset of ACs targeting IPL s1 and s3a (**Fig.1A, Supp. Fig. 1**). Using genetic tools and iterative immunostaining, we dissected Cdh4-RGCs into 7 known and 2 predicted RGC types (**Fig.2**). Two-photon calcium imaging showed that Cdh4-RGCs detected various features of dark visual stimuli and often showed preferences for stimulus color (**Fig.3**). Loss of Cdh4 leads most of these RGCs to grow dendrites into inappropriate lamina which disrupts their synaptic inputs and impairs their feature selectivity (**Fig.4,5**). Finally, we over-expressed Cdh4 in retinal interneurons which retargeted their neurites into sublamina 1-3a, likely through homophillic adhesion (**Fig.6**). Taken together, these results show a novel role for Cdh4 in the layer-targeting of nearly a third of all RGCs. Our results also define a small set of Cdhs that could encode positional information for the remaining IPL layers, suggesting a Cdh-based adhesive blueprint for layered circuit assembly in the retina (**Fig.6G**).

### A family of Cdh4 RGCs

We dissected Cdh4-RGCs into 7 known types, and 2 predicted novel types by applying a novel iterative immunostaining method called 4i^26,27^. Current taxonomies of RGCs are converging on ∼50 different mouse RGC types^4,9,10,33^. For example, a recent study matched transcriptomic and functional signatures for 42 RGC types using patch-seq^33^. Our Cdh4-RGCs are likely encompassed by this recent classification, but due to a lack of antibody markers we could not fully register our results to theirs nor fully purify our functional clustering (**Fig.3**, **Supp. Fig. 4**). Only traces within clusters 1 & 4-6 showed a selective resemblance to the mean response of their cluster; traces in clusters 2-3 also showed similarity to each other’s cluster mean. A potential advantage of the 4i method used here is that the antibodies can be stripped, and tissue stored indefinitely. Re-staining our tissue with type-specific antibody markers in the future would refine our cluster definitions and potentially separate the novel T- and F-RGC types (labelled as C9 and C32 in two recent RGC atlases^9,33^).

### Cdh4 is a laminar targeting system that directs assembly of OFF retinal circuits

We wanted to test the idea that Cdhs are generalized layer-positioning systems for retinal neurons. While we discovered new layer-specific associations for four Cdh isoforms, we focused on Cdh4. This is because we know relatively little about the function of this protein, even though Cdh4 was discovered in the chick retina (and originally named Retina-cadherin) over 30 years ago^11^. Follow-up work showed a similar Cdh4 abundance in zebrafish and mice and showed that Cdh4 labels subsets of neurons in the upper and lower INL along with many cells in the GCL^12–15^. These studies did not, however, identify the Cdh4-positive neurons or examine the function of Cdh4.

Prior work showed that Cdh4 binds strongly to itself across cell-cell junctions and is not believed to form strong heterophilic interactions in trans with other Cdhs^11,15,34–36^. Our electroporation experiments provide significance for this homophilic binding in vivo (**Fig.6**). We saw enrichment of Cdh4-overexpressing neurites in sublaminae 1 and 3a in a control background where the dendrites of many Cdh4-RGCs reside (**Fig. 6A&C**). The same Cdh4-overexpressing neurites in Cdh4-KOs were more evenly distributed (**Fig. 6B&D).** This result is consistent with the targeting deficits we saw in most Cdh4-RGC types in Cdh4- KOs, and strongly suggests that the laminar targeting effects we observed result from homophillic adhesion between neurites mediated by Cdh4.

It is important to note that our RGC targeting deficits arose without changes to the targeting of v- ooDSGCs that require Cdh6/10 for laminar targeting^6^ (**Supp.** Fig. 4C-D) and did not cause gross structural alterations to the IPL (**Supp.** Fig. 5A-F). Taken together, these results strongly support the idea that homophilic Cdh4 binding is leveraged to direct many OFF-RGC dendrites to appropriate sublamina.

An important aspect of our targeting model is that it requires a Cdh4-expressing target to be present in sublaminae 1 and 3a to act as a scaffold for actively targeting neurites. Such a neuron might express Cdh4 only to act as a scaffold but employ different methods to reach this pair of sublaminae. One possibility is that the Cdh4-ACs (**Supp. Fig. 1A-D**), could play this role. Indeed, our earlier work on starburst amacrines suggests that these neurons express Cdh6 to recruit the Cdh6/10+ve neurites of v- ooDSGCs^5^ but do not themselves require Cdh6 to find their IPL sublayers. Yet another possibility is that this scaffold cell could be one of the Cdh4-RGCs we have studied here, such as the FminiOFF, which showed attenuated phenotypes in Cdh4-KOs. More work will be needed to determine the identity of this scaffold cell.

### A combinatorial Cdh code for laminar targeting

Several factors are now known to guide the IPL ingrowth of developing dendrites and axons^37–40^. Layer targeting follows this initial ingrowth and we previously showed critical roles for Cdh superfamily members in this process for some BC and RGC types forming the retinal DS circuit^5,6^. In these studies, starburst amacrine cells expressed Cdh6 to recruit the Cdh6+ve dendrites of DSGCs to sublaminae 2 and 4. BC subsets expressed Cdh8 or Cdh9, which is required to target their axons to S2 and S4, respectively, and form synapses on ooDSGCs. The use of Cdhs in DS circuit members could have been viewed as a special adaptation to ensure development of the direction-selective circuit. Indeed, laminae containing DS circuitry are present in most vertebrates because their signals are vital to stabilize the eye^3,41,42^, and experimental or natural defects in retinal DS cause eye stabilization disorders, even in humans^42–44^. However, our results with Cdh4 and OFF-RGCs support a broader role for the Cdh superfamily in layer-targeting.

Cdh4 laminar targeting differs from our prior work in two ways. First, Cdh4 directs neurites via homophillic adhesion whereas Cdhs8&9 directed neurites with heterophilic adhesion. Second, Cdh4 directed neurites to a pair of sublaminae (s1-s3a), whereas Cdh8&9 assigned neurites to unique sublaminae. Why is this the case? Interestingly, we showed that Cdh8 or Cdh9 null BCs also target a pair of sublaminae (s2&s4) and that Cdh8 and Cdh9 assign them to one of these locations^5^ . Could this observation and our current results on Cdh4 targeting to s1-s3a reflect a two-step layer targeting governed by Cdhs? If true, then Cdh4-RGCs would need an additional factor (possibly another Cdh) to choose between s1 or s3a (**Fig. 6H**). One hint may come from our prior work which showed dendritic growth deficits in J-RGCs lacking Cdh8 or Sorcs3^6^. Misexpression studies using Cdh4 and Cdh8 could shed more light on this issue.

Finally, our transcriptomic analysis reveals layer specific associations for 3 other Cdhs (7,13, &12; **Fig 6G**). We validated layer-specific expression for Cdh13 but have not yet investigated roles of these 3 Cdhs. Nonetheless, our analysis Cdh 4, 6 and 8-10 argue for the existence of a compact Cdh-based layer-blueprint for most RGCs in the retina. Such results could also open studies into the well-known, but poorly studied, layered Cdh patterns at higher levels of the visual pathway such as the cortex^45^.

## Acknowledgements

We thank J. Sanes., E. Cooper, K. Cha, J. Lehnert, J. Forestell, J. Lefebvre, and X. Ma for helpful comments on this manuscript. We thank A. Saghatelyan for sharing the protocol for 4i in thick tissue. We thank A. Alie and J. Orlowski for help with growing electroporation constructs. We thank A.Davidova and R.Sharif for gifting us HEK cells. We thank MPBC for the loan of the 920nm fixed fiber laser used for two-photon imaging.

## Methods

### Animals

Animals were used in accordance with the rules and regulations established by the Canadian Council on Animal Care and protocols were approved by the Animal Care Committee at McGill University. Male and female Cdh4-CreER (Cdh4^Ce^), and Ai27 mice aged 0–100 days old were used in this study. Details about the generation of these lines can be found in previous studies^16,46^. Rosa26-LSL-ChR2-tdTomato mice were obtained from the Jackson Laboratory (AID27, Jackson Labs, RRID:IMSR_JAX:012567) and crossed with Cdh4-CreER mice for some anatomical experiments. A similar cross for anatomical studies involved Cdh4-CreER, Thy1-Stop15-YFP, TYW3, or HB9-GFP mice (both lines were a gift from Dr. J. Sanes). KCNG4-Cre and Vglut2-Cre mice were obtained from the Jackson Laboratory (KCNG4-Cre, Jackson Labs, RRID: IMSR_JAX: 029414; Vglut2-ires-cre, Jackson Labs, RRID:IMSR_JAX:016963). CD1 mice were obtained from Charles River Laboratories.

### Viruses

AAV9 CAG-flex-GCaMP6f, and AAV9 ef1a-flex-Tdtomato viral vectors were purchased from the Canadian Neurophotonics Platform Viral Vector Core Facility (RRID:https://identifiers.org/RRID/RRID:SCR_016477) and AAVrg-flex-Tdtomato was purchased from Addgene (viral prep: 28306-AAVrg). Retrogradely infecting AAVs were used for brain injections while AAV9 was used for intravitreal injections.

### Injections

AAVs were injected either intraocularly or intracranially to label Cdh4-RGCs using previously described methods^23^. For intracranial injections, mice were anesthetized using isoflurane (2.5% in O2), given a combination of subcutaneous carprofen and local bupivacaine/lidocaine mix for analgesia, transferred to a stereotaxic apparatus, and a small craniotomy (<1 mm) made in the appropriate location on the skull using a dental drill. Next, a Neuros syringe (65460–03, Hamilton, Reno, NV) filled with virus was lowered into either the LGN (2.15 mm posterior from bregma, 2.27 mm lateral from the midline and 2.75 mm below the pia) using a stereotaxic manipulator. A microsyringe pump (UMP3-4, World Precision Instrument, Sarasota, FL) was used to infuse 400 nL of virus (15 nL/s) bilaterally in dLGN and the bolus allowed to equilibrate for 8 min before removing the needle. For intraocular injections, mice were anesthetized as above, given carprofen as analgesic, and 0.3 μL of virus injected via an incision posterior to the ora serrata using a bevelled Hamilton syringe (7803-05, 7634-01, Hamilton). Mice were given at least 2 weeks to recover before experimental use.

### Tamoxifen

Tamoxifen (TMX) (MillliporeSigma, Cat#T5648) was dissolved in anhydrous ethanol at 200 mg/mL, diluted in sunflower oil to 10 mg/mL, sonicated at 40°C until dissolved, and stored at –20°C. Prior to injection, TMX aliquots were heated to 37°C and delivered intraperitoneally at ∼1 g/50 g body weight to Cdh4-CreER ∼P30 mice. The dose was repeated twice over 2 days and mice were used between 2-4 weeks following treatment. For sparse labeling we use a single dose of TMX at ∼.05 g/50g body weight.

### Histology

Mice were euthanized by isoflurane overdose and transcardially perfused with chilled PBS followed by 4% (w/v) paraformaldehyde (PFA) in PBS and enucleated. Eye cups were fixed for an additional 45 min in chilled 4% (w/v) PFA. For whole-mount stains, tissue was incubated in primary antibodies for 7 days at 4°C and incubated in secondary antibodies overnight at 4°C following a series of washes. For cross-sections, post-fixed retinae were sunk in 40% (w/v) sucrose/PBS, transferred to embedding agent (OCT, Tissue-Plus, Fisher Scientific), flash frozen in 2-methylbutane at –45°C, and then sectioned onto slides at 30 μm thickness on a cryostat. For immunostaining, slides were first washed in PBS, blocked with blocking solution (4% normal donkey serum, 0.4% Triton-X-100 in PBS) for 2 hr at room temperature (RT) and incubated overnight with primary antibodies at 4°C. Tissue was then washed with PBS and incubated in secondary antibodies for 2 hr at room temperature prior to the final wash and tissue mounting. HEK cells were fixed for 10 min with 4% PFA, rinsed with PBS and immunostained as described for cross-sections. For nuclear staining NucBlue (R37606, Fisher Scientific) was added during rinsing after secondary antibody incubation.

### 4i immunostaining

Retina stains following two-photon calcium imaging and for cell type lamination were performed using a modified version of 4i immunostaining^26,27^. Briefly, retinas were fixed in pre-chilled 4% PFA for 45min and rinsed with PBS, followed by 15 min in pre-chilled methanol and 15min in pre-chilled acetone at 4°C. After rinsing, the tissue was blocked with 4% blocking solution for 2 hours at RT; incubated with primary antibodies overnight at 4°C and in secondary antibodies for 3 hours at RT; mounted and imaged. We used an imaging buffer (0.7M N-Acetyl-Cysteine in 0.2M phosphate buffer, pH 7.4) as mounting medium to prevent antibody crosslinking with the tissue. After imaging, we proceeded to elute antibodies from the tissue by rinsing it for 10min in ddH2O at RT and applying an elution buffer (0.5M L-Glycine, 3M Urea, 3M Guanidine Hydrochloride, 70mM TCEP-HCl (TCEP), pH 2.5) at RT. After elution we re-stained using the same procedure as many times as necessary. In retinas used for calcium imaging, lectin was included during secondary antibody incubation to stain blood vessels for registration.

### Antibodies and stains

Antibodies used were as follows: rabbit anti-DsRed (1:1000, Clontech Laboratories; RRID:AB_10013483); guinea pig anti-DsRed (1:500, Synaptic Systems, 390 004); chicken anti-GFP (1:1000, Abcam, Cambridge, UK; RRID:AB_300798); rat anti-Cdh4 (1:100 DSHB, RRID: AB_528110); mouse anti-Cdh13 (1:500 Santa Cruz Biotechnology, RRID:AB_10612090); goat anti-VAChT (1:1500, MilliporeSigma, Darmstadt, DE; RRID:https://identifiers.org/RRID/RRID:AB_2630394); guinea pig anti-RBPMS (1:100, Phosphosolutions, Aurora, CO; RRID:https://identifiers.org/RRID/RRID:AB_2492226); mouse anti-Ap2-α (1:100, clone 3b5 from Developmental Studies Hybridoma Bank, Iowa City, IA); mouse anti-Chx10 (1:300, Santa Cruz Biotechnology; RRID:https://identifiers.org/RRID/RRID:AB_2216006); mouse anti-Brn3c (1:250, Santa Cruz Biotechnology, Dallas, TX; RRID:https://identifiers.org/RRID/RRID:AB_2167543); rabbit anti-Tbr1 (1:1500 Abcam;, RRID:AB_22002); rabbit anti-Tbr1 (1:1000 Abcam ab183032); goat anti-osteopontin (1:1000, R&D Systems, Minneapolis, MN; RRID:https://identifiers.org/RRID/RRID:AB_2194992); goat anti-Foxp2 (1:1500 Abcam; RRID:AB_1268914); rabbit anti-PKCα (1:1000, Sigma Aldrich, P4334); rabbit anti-calbindin (1:10000, Swant, CB38); guinea pig anti-Vglut3 (1:2000, Millipore Sigma, AB_5421).and. Secondary antibodies were conjugated to Alexa Fluor 405 (Abcam; RRID: https://identifiers.org/RRID/RRID:AB_2715515 or Jackson ImmunoResearch Labs; RRID:https://identifiers.org/RRID/RRID:AB_2340427), Alexa Fluor 488 (Cedarlane, Ontario, CA; RRID:https://identifiers.org/RRID/RRID:AB_2340375), FITC (MilliporeSigma; RRID:https://identifiers.org/RRID/RRID:AB_92588), Cy3 (MilliporeSigma; RRID:https://identifiers.org/RRID/RRID:AB_92588, RRID:https://identifiers.org/RRID/RRID:AB_92570, or Jackson ImmunoResearch, West Grove, PA; RRID:https://identifiers.org/RRID/RRID:AB_2340460, RRID:https://identifiers.org/RRID/RRID:AB_2340694) or Alexa Fluor 647 (MilliporeSigma; RRID:https://identifiers.org/RRID/RRID:AB_2687879). Isolectin (Fisher Scientific, Waltham, MA; RRID:https://identifiers.org/RRID/RRID:SCR_014365) was incubated along with the secondary antibodies when applicable. NucBlue (Fisher Scientific, R37606) was incubated after secondary antibodies when applicable.

### Confocal Imaging and analysis

Images of stained tissue were acquired on an Olympus FV1000 confocal laser-scanning microscope or a Zeiss LSM-710 inverted confocal microscope (Advanced BioImaging Facility, McGill University) at a resolution of 512 × 512 or 1024 × 1024 pixels with step sizes ranging from 0.3 μm to 8 μm.

To obtain mean IPL projection depth of each Cdh4-type, we took linescans of pixel intensity across the IPL from images stained with antibodies to VAChT (sublamina 2 and 4) or reporter, normalized these signals to the maximum intensity, and averaged these traces across each condition. IPL depth is expressed as a percent and sublaminae judged by the position of the peak intensities in the VAChT channel. We applied the straighten transform (ImageJ) to the VAChT bands on curved sections and applied the same transform to reporter channels prior to linescan procedure.

### Calcium imaging

Calcium imaging was performed as previously described^18,23^. Briefly, mice were dark adapted for at least 2 hr, euthanized, and then retinae rapidly dissected under infrared illumination into oxygenated (95% O2; 5% CO2) Ames solution (MilliporeSigma, A1420). Next, retinae were mounted onto a filter paper (MilliporeSigma, HABG01300) with the RGC layer facing up, placed in a recording chamber, mounted on the stage of a custom-built two-photon microscope, and perfused with oxygenated Ames solution warmed to 32–34°C. Responses of GCaMP6f+ RGCs to visual stimuli delivered through the objective were imaged at 920 nm (MPB Femto fixed 920nm laser) and collected at an imaging rate of 45 Hz. Each image plane (323.74 x 323.74um μm) of the movie contained GCaMP fluorescence, SR101 fluorescence, stage coordinates, and visual stimulus synchronization pulses to permit offline analysis. A few microliters of sulphorhodamine 101 (SR101, 2 mg/mL, MilliporeSigma, S7635) was added to the recording chamber to label blood vessels and a map of the main blood vessels emanating from the optic disk acquired for post hoc image registration^23^. Data acquisition was performed with ScanImage synced to the visual stim using a photodiode sync pulse and handshake pulse using custom made code in Matlab 2023. Following recording, retinas were fixed, immunostained and imaged as described above.

### Data pre-processing

Calcium responses were aligned with stimuli using custom made Matlab code. Regions of interest (ROIs) were defined using CellPose^47^ and Matlab (R2020b) was used to extract Ca2+ traces for each ROI. The signals were smooth to remove noise with moving mean of 10 and expressed as ΔF/F, where the mean of the peristimulus baseline was used to calculate the difference. Then, a quality index representing signal-to-noise ratio was calculated to filter out not reliable cells as described before (Bade et al., 2016) using moving bar traces, only cells with a QI > .45 were used for further analysis.

Several descriptive statistics were calculated for RGC calcium responses: ON-OFF index: (OOi) was calculated as the vector representation in 2D of the ON and OFF responses to a full-field flash.

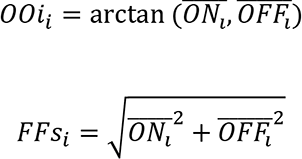

Where the magnitude of the vector (FFs) represents how selective the cell is for On vs Off, with higher values indicating high selectivity; and the angle (OOi) represents the On-Off preference (ON-OFF, 0:90; ON, -180:0, OFF, 90-180).

<u>Frequency selectivity and preference</u>

Frequency selectivity (FQ-s) was calculated as the vector sum of responses to each of the 21 different frequencies (f) shown in the fq-chirp stimulus. Each cell response was fitted to an envelope, and the maximum response to each frequency was assigned as a value for one dimension of a frequency vector (F). The magnitude represents how selective the cell is to a specific frequency, where:

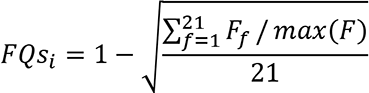

FQ-s index ranges from 0 to 0.7818 (0.7818 = cell is frequency selective; 0 = cell is not frequency selective). The preferred frequency (FQ-p) for each cell was defined as the f corresponding to the max(F).

<u>Contrast selectivity and preference</u>

Contrast selectivity (CN-s) was calculated using the same method as for frequency with a vector sum across 16 contrast changes (C). CN-s index ranges from 0 to 0.75 (0.75= cell is contrast selective; 0 = cell is not contrast selective). The preferred contrast (CN-p) for each cell was defined as the contrast percentage corresponding to the max(C).

Green-Blue index: (GBi) was calculated using the maximum Off or On response of each cell to the green (G) and blue (B) full-field flashes:

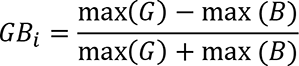

The GBi ranged from -1 to 1 (-1 = cell responds only to blue; 0 = cell responds equally to both colors, 1 = cell only responds to green) For full-field, fq-chirp, cn-chirp and full-field color the amplitude was defined as the area under the curve (AUC) for each response. For experiments used to characterize direction selective RGCs in Cdh4-KOs an index (DSI) was computed according to standard methods of circular variance^23,30^.

### PCA and Clustering

Calcium responses were down sampled to 3Hz for feature extraction and clustering. Principal component analysis was implemented in R (version 4.1.0) to extract features from our 6 different stimuli. A total of 14 features were further used for clustering using a diagonal, varying volume, and shape gaussian mixture model (GMM). This model was evaluated using the Bayesian information criterion to select the optimal number of clusters. This method resulted in 7 clusters but one of them was removed after all control and Cdh4-KO were classified giving it contained cells with high stimulus contamination These clusters were refined using the silhouette coefficient to reassign observations to new clusters. If the silhouette coefficient of a cell was negative the observation was reassigned to the cluster that showed the minimal average dissimilarity. This process was repeated iteratively 22 times until the number of positive silhouette values for all observations reached a plateau and the GMM was modified accordingly.

To assign data from Cdh4-KO RGCs into the 7 clusters defined using Cdh4-Het controls, first, calcium responses from Cdh4-KO were projected into the new PC space using the eigenvectors obtained from controls. Then, using the optimal GMM from the control data set, Cdh4-KO cells were evaluated for their probability to belong to each gaussian and finally assigned to the cluster with the highest probability. Additionally, a new data set from Cdh4-Het control was assigned using this same method to add cell-type markers to each cluster. A uniform manifold approximation and projection (UMAP) was used for visualization.

To check the consistency of the clustering we compare the responses for all stimuli of each cell within each group to the average cluster response of the 7 clusters. To do this, we normalized each cell trace by its Euclidean length and computed the cosine similarity of each cell to the normalized mean response of each cluster.

Retinas were 4i immunostained after two-photon calcium imaging and image registration was performed as described previously^23^. Briefly, the average of each recorded field was stitched into a single image using custom MATLAB scripts, while a confocal image was acquired for the same field after 4i staining. Following scaling, rotation, and translation the two images were overlayed to match sulforhodamine and lectin blood vessel patterns. Next, fine blood vessel morphology was used as inputs for landmark correspondence in Fiji to register each two-photon confocal field with 4i confocal images. Following registration, each ROI obtained with Cellpose (see above) was semi-manually assigned markers using custom MATLAB scripts. For marker assignment, only cells with a group similarity above average of its assigned cluster were used.

### In Vivo Electroporation

A full-length cdh4 C-OFPSpark tag cDNA was obtained from SinoBiological (MG51158-ACR). For expression, the constructs were transferred to a pCAGGS expression vector either alone or with aa Cre-dependent “flex-switch” by Epoch Life Sciences. For control experiments we used a pCAG-mCherry plasmid.

*In vivo* electroporation was performed as described previously^48^. Briefly, neonatal mice (P0-P1) were injected using a glass pipette or a Hamilton syringe with a 32-gauge blunt-ended needle into the subretinal space with plasmid (1-.5ul of ≈1mg/mL) in PBS containing a 10% of fast green. After DNA injection, current pulses (Unipolar, 80V, 50ms, 5 pulses, 1Hz) were applied on the head using electrode tweezers (Protech Internationa; CUY650-P5) and a pulse generator ECM830 (BTX-Harvard apparatus; 45-0052INT). Retinas were collected at P21 and dissected for immunohistochemestry.

### Cell culture

HEK293 cells were transfected with the pCMV3-C4-OFPSpark plasmid using lipofectactamine in 35mm plates. After 24 hours of transfection, cells were fixed using 4% PFA for 10min at RT and stained.

### Electrophysiology

Retinae for electrophysiological recordings were prepared as previously described^23,25,49^. For cell-attached recordings, the patch electrodes (4–5 MΩ) were filled with Ringers solution. For whole-cell recordings, patch electrodes (5–7 MΩ) were filled with an internal solution containing 112 mM Cs methanosulfate, 10 mM NaAc, 0.2 mM CaCl2, 1 mM MgCl2, 10 mM EGTA, 8 mM CsCl, and 10 mM HEPES (pH 7.4). In both cell-attached and whole-cell recordings, fluorescein 3000 MW dextran (Thermo Scientific, D7156) was added to make the electrode visible under two-photon illumination. For whole-cell recordings, internal solution was supplemented with 5 mM QX314 Bromide. Excitatory currents and inhibitory currents were isolated by adjusting the holding potential to match reversal potentials for excitation (+15 mV) and inhibition (ECl ∼ –60 mV). Signals from loose-patch and whole-cell recordings were acquired with a MultiClamp 700B amplifier (Molecular Devices) and digitized at 20 kHz using custom software written in LabView. For spikes, the MultiClamp was put into I = 0 mode and Bessel filter set at 1 kHz. For currents, the MultiClamp was put in VC mode and Bessel filter set at 3 kHz. Analysis of electrophysiological signals was performed in MATLAB (Simulink) as follows. Briefly, action potentials were detected in loose patch recordings using the *peakfinder* function and binned (50 ms) over the entire length of the trial; firing rate histograms for each trial were then averaged and subjected to further processing based on each stimulus. For whole-cell currents, trials were averaged, peak amplitude measured, and integral were computed using the *trapz* function across each stimulus epoch.

### Visual Stimuli

A DLP light crafter (Texas Instruments, Dallas, TX) was used to project dichomatic (405nm, 520 nm) visual stimuli through a custom lens assembly that steered stimulus patterns into the back of a 16× objective^18,23^. All visual stimuli were written in MATLAB using the psychophysics toolbox and displayed with a background intensity set to 1 × 10^4^ R*/rod/s. Custom electronics were made to synchronize the projector LED to the scan retrace of the two-photon microscope^50^.

For calcium imaging experiments, visual stimuli were centered around the microscope field of view. Moving bar stimulus consisted of a bright bar moving along its long axis in one of eight directions. The bar was 300 μm wide, 1500 μm long moving at 1000 μm/s (data not shown). Full-field stimulus was presented with increment and decrement of light that lasted 2s. Freqeuncy chirp consisted of a full contrast sinusoidal intensity modulation from 1 to 16Hz. Contrast chirp consisted of a 4Hz synodal with 8-bit contrast increase per cycle. Color stimuli were delivered using full flashes with only UV or green LEDs that lasted 3s. For all full-field stimuli a 2000μm diameter circle in a gray background was used. For electrophysiological experiments, the cell-receptive field center was identified using a grid of flashing spots and a small user-controlled probe and the location with the highest response assigned as the center for all subsequent stimuli. All stimuli were preceded by a gray background.

### RNA seq Data Analysis

scRNA sequencing data was retrieved from the publicly available GEO database: RGCs, GSE137400; ACs, GSE149715; and BCs, GSE81905^7–9^. To identify Cdhs differentially expressed in each individual dataset, a differential expression analysis was performed using Seurat (version 4.0.3) in R Sequencing data plots were generated with the ggplot2 plot package (version 3.3.6) in R.

Dendrogram of Cdh RGC expression was built using hierarchical clustering in R (hclust). Briefly, this method uses a set of dissimilarities for iteratively cluster assignment of RGC subtypes by implementing the Lance-Williams update formula.

### Statistics

No statistical method was used to predetermine sample size. Statistical comparisons between Cdh4-KO and Cdh4-Het electrophysiological, morphological, and calcium imaging data were performed using t-test in MATLAB and R. Group similarity significance was tested using ANOVA and Tukey Honest Significant Differences with the tukey_hsd() function in the rstatix package in R.

## Supplemental Figures

**Supplemental Figure 1.**
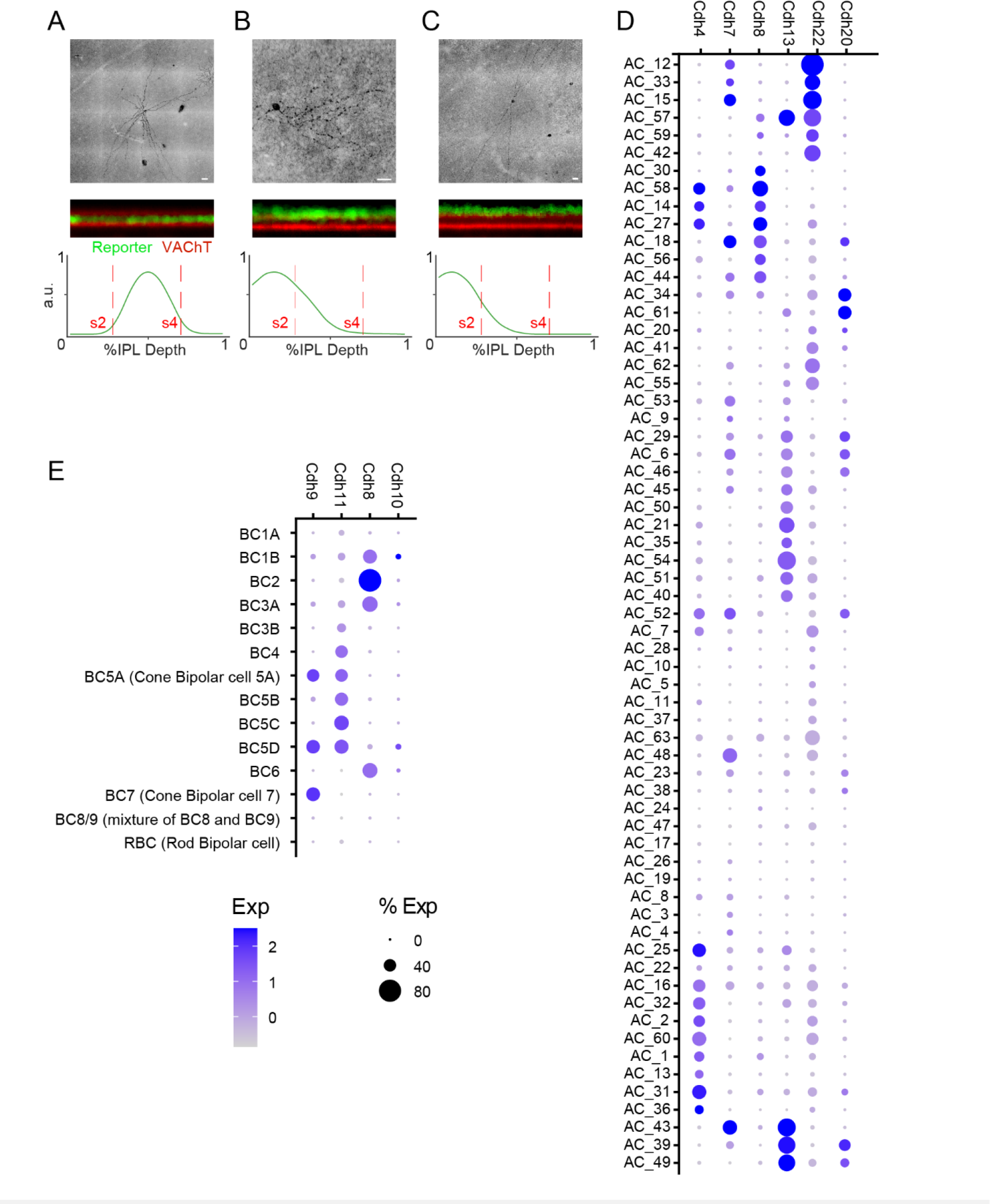
Cdh expression in interneurons. A-C. Cdh4+ve ACs sparsely labelled with Cre-dependent reporter in retinal wholemounts counterstained with antibodies against VAChT (top), in rotated z-stacks to resolve laminar position (middle), and linescans through the examples shown in middle. **D.** Dotplot showing expression of Cdhs4,6,8,13,20, and 22 in amacrine cells . **E.** Dotplot showing expression of Cdhs9,11,8, and 10, in bipolar subtypes.

**Supplemental Figure 2.**
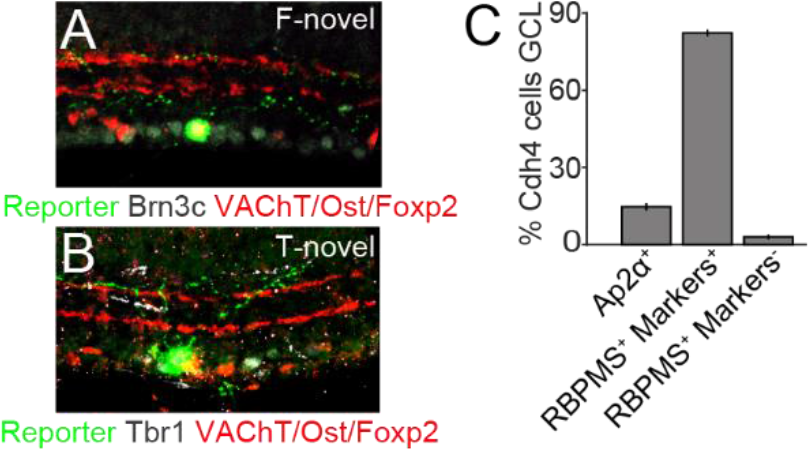
Novel Cdh4 RGC cell types. A. Retinal cross-section from a Cdh4-Het x tdtomato stained showing a foxp2+ve RGC that are YFP-, Osteopontin- and Brn3c+. **B.** Retinal cross-section from a Cdh4-Het x tdtomato mouse showing a Tbr1+ve RGC that is Osteopontin- and Foxp2-. **C.** Percentage of GCL cells (±SEM) in Cdh4-Het retinas labeled with RBPMS and either Tbr1/Brn3c/Foxp2/Ost (RBPMS+ Markers+), with only RBPMS (RBPMS+ Markers-), or with Ap2α (Ap2α+).

**Supplemental Figure 3.**
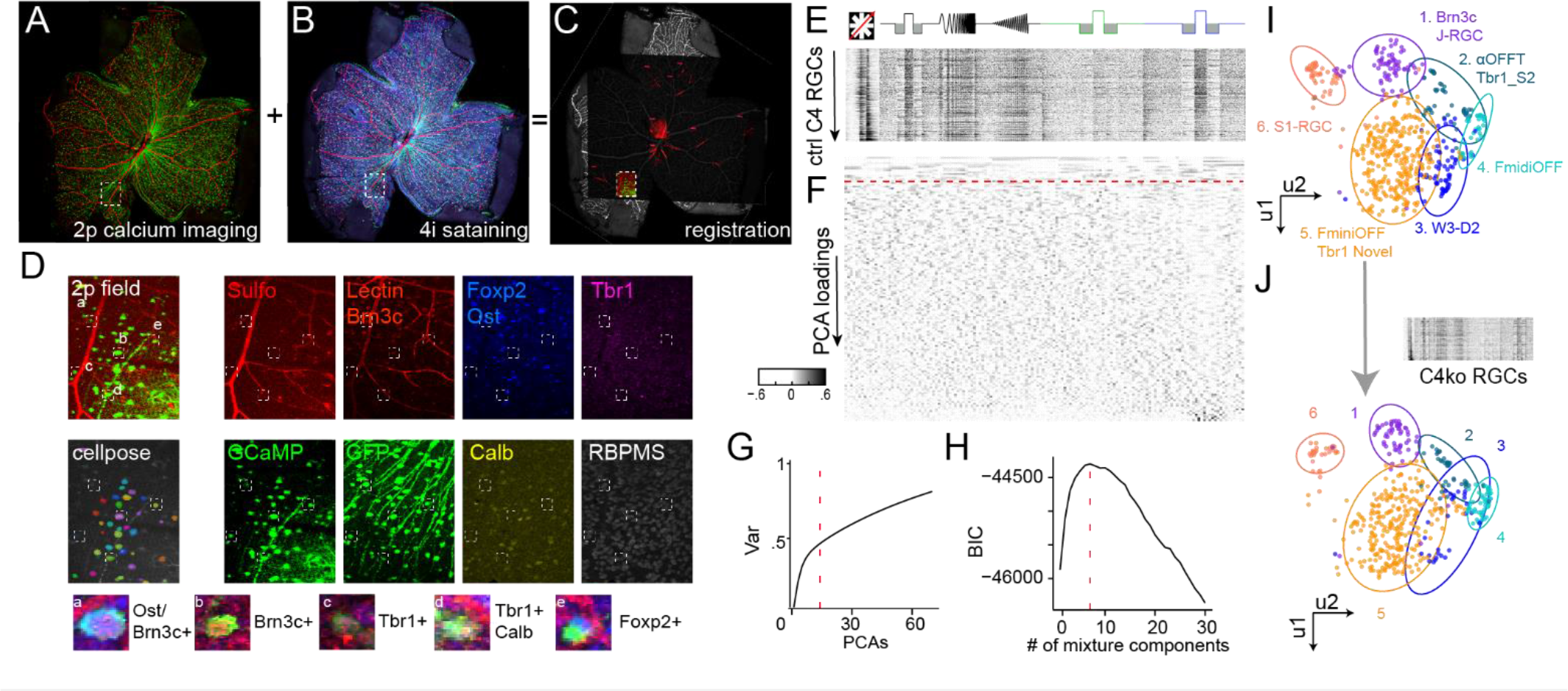
Marker registration and clustering. A-C. Whole-mount retina labeled for blood vessels and GCaMP6f+ve cells used during two-photon imaging (A) were registered to cell markers acquired with 4i staining (B,C). **D**. Example of a two-photon field with ROIs found using cellpose and registered to low-power images with cell markers. Bottom, insets for example cell types found using this method. **E.** Cdh4-Het response matrix to different visual stimuli obtained from cellpose ROIs. **F,G**. PC loadings (F) from responses in E and the variance they explain (G). Dashed line indicates the cutoff for PCs selected for dimension reduction (PCs = 14). **H,I.** PCs were used to cluster Cdh4 cells with the best GMM selected based on the information contained in its component number (H, components = 7) and visualized with UMAP (I). Cluster 7 was removed because it contained stimulus contaminated cells. **J.** Cdh4-KO response matrix (top) is cluster using the PC loadings (F) and GMM (I) from Cdh4-Het cells.

**Supplemental Figure 4.**
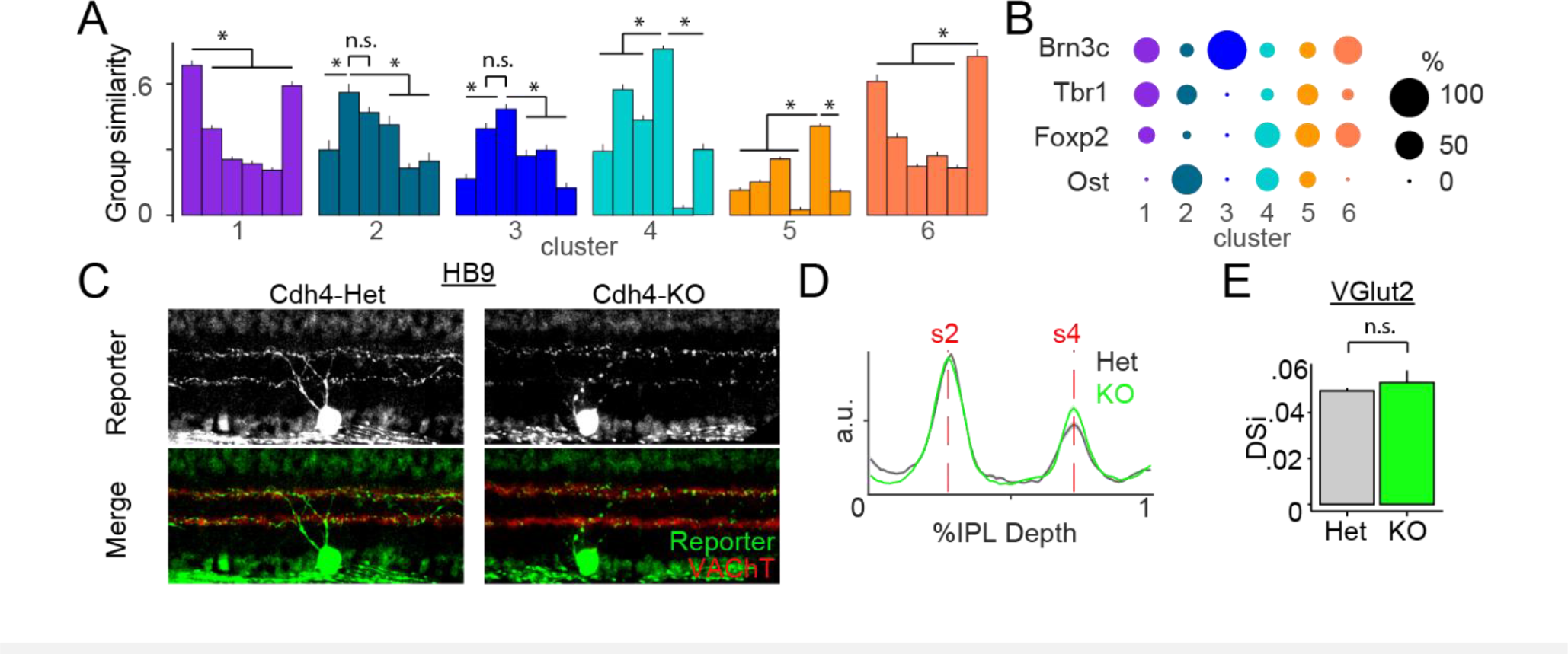
Molecular markers define cluster identity and Cdh4 loss does not impact morphology of Cdh4-ve RGCs. A. Cosine similarity computed for RGCs in each Cdh4-Het cluster (±SEM). * = P < 0.05 **B.** Marker expression for each Cdh4-Het cluster. Cluster 2 and 3 do not show significant difference (n.s) between their group similarities (A) but makers set apart these two clusters (B) indicating strong resemblance in their visual responses. **C-D.** Direction-selective RGCs labelled in Cdh4-Het x HB9(C, left) and Cdh4-KO x HB9 (C, right) retinal cross-sections, and mean line scans (±SEM) taken through their dendritic arbors (D). Loss of Cdh4 does not impair laminar targeting of the Cdh6/10-expressing neurons. n ≥ 7 for each condition for a total of 5 mice **E.** Direction selective index from Ai32-G6F x VGlut2Ce retina in Cdh4-Het and Cdh4-KO conditions.

**Supplemental Figure 5.**
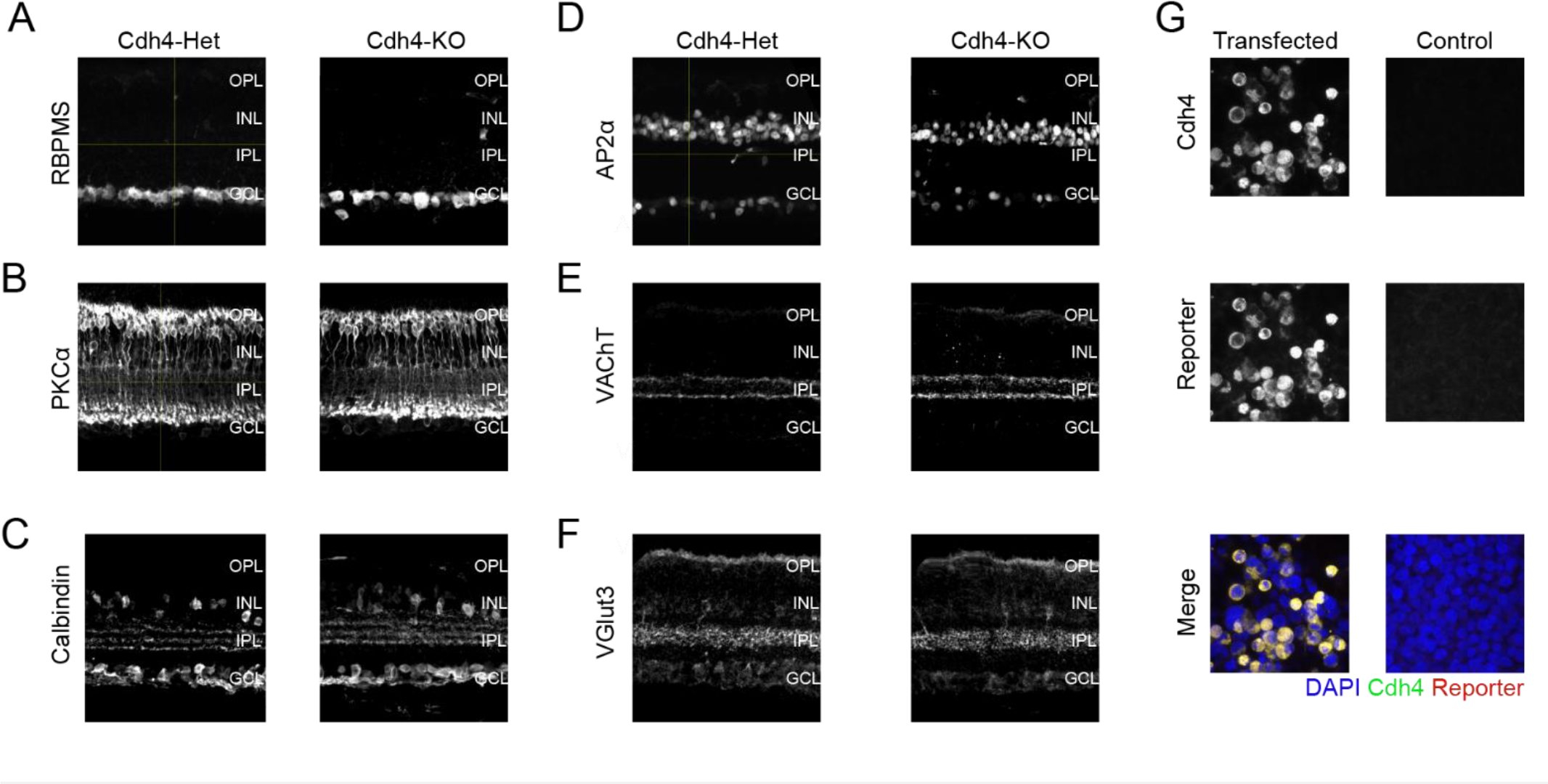
Cdh4 loss not grossly alter to retinal structure A-F. Retinal cross sections taken from Cdh4-Het and Cdh4-KO mice stained with antibodies against RBPMS (A), PKCα (B), calbindin (C), Ap2α (D), VAChT (E) and Vglut3 (F). **G**. HEK cells transfected with constructs encoding Cdh4-OFP and counterstained with anti-Cdh4 antibodies (left). Untransfected controls are also shown (right).

